# Shear rheology of methyl cellulose based solutions for cell mechanical measurements at high shear rates

**DOI:** 10.1101/2022.11.18.517048

**Authors:** Beyza Büyükurgancı, Santanu Kumar Basu, Markus Neuner, Jochen Guck, Andreas Wierschem, Felix Reichel

**Affiliations:** Max Planck Institute for the Science of Light and Max-Planck-Zentrum für Physik und Medizin, Erlangen, Germany; Institute of Fluid Mechanics, Friedrich-Alexander-Universität Erlangen-Nürnberg (FAU), Erlangen, Germany; Chair of Biological Optomechanics, Department of Physics, Friedrich-Alexander-Universität Erlangen-Nürnberg (FAU), Erlangen, Germany

## Abstract

Methyl cellulose (MC) is a widely used material in various microfluidic applications in biology. Due to its biocompatibility, it has become a popular crowding agent for microfluidic cell deformability measurements, which usually operate at high shear rates (> 10,000 s^−1^). However, a full rheological characterization of methyl cellulose solutions under these conditions has not yet been reported. With this study, we provide a full shear-rheological description for solutions of up to 1% MC dissolved in phosphate-buffered saline (PBS) that are commonly used in real-time deformability cytometry (RT-DC). We characterized three different MC-PBS solutions used for cell mechanical measurements in RT-DC with three different shear rheometer setups to cover a range of shear rates from 0.1 - 150,000 s^−1^. We report viscosities and normal stress differences in this regime. Viscosity functions can be well described using a Carreau-Yasuda model. Furthermore, we present the temperature dependency of shear viscosity and first normal stress difference of these solutions. Our results show that methyl cellulose solutions behave like power-law liquids in viscosity and first normal stress difference at shear rates between 5,000 - 150,000 s^−1^. We construct a general viscosity equation for each MC solution at a certain shear rate and temperature. Furthermore, we investigated how MC concentration influences the rheology of the solutions and found the entanglement concentration at around 0.64 w/w%. Our results help to better understand the viscoelastic behavior of MC solutions, which can now be considered when modelling stresses in microfluidic channels.

## 1 Introduction

Methyl cellulose (MC) is one of the most important cellulose ethers, widely used in food processing^1–3^, pharmaceutical productions^4,5^ and cosmetics.^6,7^ Because of its biocompatibility, it is a popular crowding agent in cell research.^8–10^ Recently, MC was employed in microfluidic cell mechanical measurements because of its ability to increase the cell carrier viscosity already at small concentrations while maintaining cell viability. An increased viscosity reduces cell sedimentation during the experiments and increases the hydrodynamic shear stresses that act on the cell surface to deform them, which is often desirable.^8,9,11,12^ As cell carrier for cell mechanical measurements in real-time deformability cytometry (RT-DC), MC is dissolved in phosphate buffered saline (PBS).^8,13,14^ For microfluidic cell mechanical measurements, it is important to understand the rheological behavior of the MC-PBS solutions over a wide range of shear and extension rates to simulate the flow conditions in the microchannels and to understand the stresses acting on the cell surface.^11,15–18^

The solutions are usually not characterized at the high shear rates present in these microchannels since conventional rheometers cannot operate under these conditions. The rheology of aqueous MC-solutions, for instance, has been measured at shear rates up to 5,000 s^−1^, where the shear-thinning regime followed a power law.^7,11,15,16^ Furthermore, the addition of salt only influenced the shear and extensional viscosity at high concentrations^16^ and the pH value did not show any remarkable influence on the rheology of the MC-solutions.^19^ The viscoelastic properties of MC-solutions were described by Amari and Nakamura^20^ who found that the polymer solutions were in an entangled regime at concentrations below 1.0%. The shear thinning behavior of the MC-PBS solutions at shear rates up to 150,000 s^−1^ was described by Herold^15^, but this study was not performed under controlled temperatures and did not report the viscoelastic behavior of the solutions.

Here, we combined the results from three shear rheometers to provide a full shear-rheological description of three MC-PBS solutions used for RT-DC experiments. We measured the shear viscosity using a concentric-cylinders (CC), cone-plate (CP) and parallel disks (PP) device to cover a large range of shear rates of 0.1 – 150,000 s^−1^. While the viscosity of these solutions was presented before^15^, we revisited these results and now provide viscosity curves at controlled temperatures. To better understand the entanglement regimes of the MC polymer solutions used in this study, we measured MC concentrations between 0.3 – 1.0 w/w% in the CP device and determined the zero viscosities to find the entanglement concentration. Our results provide a full shear-rheological description of MC-PBS solutions. These results can be used to model the flow behavior in any microfluidic application, e.g., to compute the stresses on cell surfaces in RT-DC. Especially the effect of the normal stress differences should not be neglected, which were, to our knowledge, not described before. These insights will help to improve our general understanding of flow systems with MC solutions.

## 2 Materials and Methods

### 2.1 Methyl cellulose solution production

Methyl cellulose (MC, 4000 cPs, Alfa Aesar 036718.22, CAS# 9004-67-5) and phosphate buffered saline (PBS) (Gibco Dulbecco 14190144) were used to prepare MC-PBS solutions. Osmolality of PBS was adjusted to values between 280-290 mosm/kg before adding MC (VAPRO Vapor Pressure Osmometer 5600, Wescor). The three solutions studied here were named 0.49% MC-PBS, 0.59% MC-PBS and 0.83% MC-PBS and were prepared as follows: To prepare 0.49% MC-PBS, 0.59% MC-PBS and 0.83% MC-PBS solutions for 1 L of PBS (1010 g), 4.90 g, 6.00 g and 8.40 g MC powder were used, respectively. The weight concentrations of the MC-PBS solutions were 0.485 (w/w%) for 0.49% MC-PBS, 0.594 (w/w%) for 0.59% MC-PBS and 0.832 (w/w%) for 0.83% MC-PBS. After adding the powder, the mixtures were kept at 4 °C under constant rotation for two days while the MC dissolved. For this, we placed a rotating mixer (RS-TR05, Phoenix Instrument) in a High-Performance Pharmacy Refrigerator (TSX Series, Thermo Fisher Scientific) and kept the bottles with the MC-PBS solution mixing inside. Then the pH value was adjusted to 7.4 (Orion Star A211 benchtop pH meter, Thermo Scientific) by addition of NaOH. In a next step, the solutions were sterile filtered using membrane filters (Merck Millipore Steritop-GP polyethersulfone (PES) membrane, 0.22 μm pore size). The buffer solutions were stored at 4 °C.

To check the influence of the MC concentration, we prepared solutions of 0.3, 0.4, 0.5, 0.6, 0.7, 0.8, 0.9 and 1.0 (w/w%) MC in PBS. These solutions were used for measurements in the CP device only. The same type of methyl cellulose and PBS were used to prepare the solutions as described above. To produce the solutions, we dissolved the MC powder in PBS and adjusted the total weight to 50 g. For example, for 1.0 (w/w%) MC-PBS we used 1.0 g of MC powder in 49.0 g of PBS. Note that these solutions were not to be used in cell measurements, thus pH value adjustment and filtration steps were not applied.

### 2.2 Shear rheometry

#### 2.2.1 Concentric cylinders (CC)

The lowest shear rates were studied with a concentric cylinder geometry on an MCR 702 Multidrive (Anton Paar GmbH, Graz, Austria). Temperature control was carried out using a water-cooled CTD 600 measuring chamber. A Searle system C-CC20/TD (diameter: 22 mm) cup and B-CC20 (diameter: 20 mm, length 30 mm, cone tip 120°) bob was used. The filling quantity and immersion depth were determined according to DIN EN ISO 3219-2 and amounted to 8.17 mL and the distance between the tip of the cone and the bottom of the beaker was 4.23 mm.

The sample was tempered to 25±0.2 °C before it was pre-sheared for 30 seconds at a shear rate of 10 s^−1^ followed by 3 min of rest. For each sample, forward and reverse measurements were carried out at shear rates between 10^−1^-10^3^ s^−1^. To minimize the impact of imperfect concentricity, the measuring time per data point of the individual shear rates usually corresponds to the duration of one complete revolution of the bob but was at least 20 s. After increasing the shear rate stepwise to the highest value, we decreased it likewise to the lowest. Here, we show the data of this backward measurement, which showed high reproducibility with repeated forward and backward measurements (variation less than 3%), while the initial forward run deviated from them by up to 10% despite pre-shearing.

Directly after the unidirectional shear measurements, an amplitude sweep followed by a frequency sweep were carried out with the same sample. In the amplitude sweep, the frequency was fixed to 1 Hz and the measurement time per data point to 20 s. The deformation was increased stepwise from 1% to 500% and vice versa. The deformation amplitude for the frequency sweep was set to 4%, which was well within the linear viscoelastic range, and the frequency was varied from 0.01 Hz to 100 Hz and back again. The measurement time per data point was between 5 - 250 s and was determined by the software (Rheocompass, Anton Paar GmbH, Graz, Austria).

#### 2.2.2 Cone-plate (CP)

Cone-plate measurements were performed using the Anton Paar MCR 502 TwinDrive rheometer (Anton Paar GmbH, Graz, Austria) combined with a PTD 180 MD Peltier element (Anton Paar GmbH, Graz, Austria) for temperature control. A CP40-0.3 (diameter: 40 mm, angle: 0.3°) cone and PP50 (diameter: 50 mm) parallel-plate (both Anton Paar GmbH, Graz, Austria) were used as upper and lower measuring system, respectively.

To avoid evaporation of the sample during the measurements, the heating chamber was equipped with a water reservoir under the lower plate and wet paper tissues were added to the walls of the chamber to create a solvent trap. Before each measurement, we performed a zero-gap detection and both upper and lower motor were calibrated. MC samples were loaded on the lower plate with a volume of 210 μl with a micropipette (Eppendorf Research® plus 1000 μl, Hamburg, Germany). The nominal gap volume of this measuring system is 90 μl but we intentionally used large overloading to take out edge effects at the sample^21^, like local gelation of the solution due to evaporation at the air-sample-interface.

When moving to the measurement position after sample loading, the lower plate was rotating to have a good volume distribution and bubble-free sample. To release possible internal stresses in the sample before measuring, a pre-shearing stage was applied at 200 s^−1^ for 30 seconds. Normal force and torque were monitored before starting the measurements without any shear, to make sure the sample is in a relaxed steady state. Once the sample was relaxed, normal force and torque were reset to zero.

For the measurement of MC solutions at room temperature (25 °C), a shear rate sweep was applied in two different shear rate intervals. The first interval started at 1 s^−1^ and ramped logarithmically up to 100 s^−1^ over 10 data points. The first data point was measured for 10 seconds, and measurement time for each data point was decreased logarithmically to 2 s at the highest shear rate of 100 s^−1^. In this shear rate range, we used the separate motor transducer mode (SMT) of the rheometer, where only the lower plate is rotating and the torque is measured from the upper plate, due to its higher sensitivity as compared to the counter rotation mode. At higher shear rates up to 25,000 s^−1^, we used the counter-rotation mode to avoid radial migration. In this range, each data point was measured for 2 seconds and 12 data points were taken in the logarithmically distributed shear rate interval. This measurement time ensures a stabilized shear stress to determine the viscosity at any shear rate. Despite these measures, radial migration restricted the maximum shear rate to 20,000 s^−1^ in the 0.49% MC-PBS samples. Only forward measurements were considered for data analysis. Measurements were also conducted in backward direction (from 25,000 s^−1^ to 1 s^−1^) to check if some amount of sample was lost during the rotation. If so, the measurement was not considered for further analysis.

Temperature dependent measurements were carried only with the cone-plate geometry and aimed only for higher shear rates. These shear rate sweeps were performed in an interval between 600 s^−1^ and 25,000 s^−1^ in counter-rotation mode. 12 data points were taken by keeping the measurement time constant at 2 seconds for each data point. Viscosity and 1^st^ normal stress differences (*N*_1_) were measured at 22, 25, 28, 31, 34 and 37 °C covering the relevant range for experiments on biological cells.

1^st^ normal stress differences of the samples in cone-plate geometry, in counter-rotation, were calculated with equation (1)^22^:

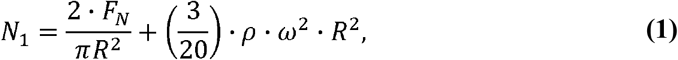

where *F_N_* is the measured normal force, *R* is the cone radius, *ρ* is the fluid density and *ω* is the angular velocity of the lower plate only. The second part of equation (1) describes effects due to inertial migration and was here adapted to fit the system in counter-rotation. Angular velocity can be defined as following for one plate:

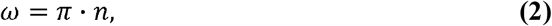

where *n* is the rotational speed.

#### 2.2.3 Parallel disks (PP)

To access shear rates beyond 10^5^ s^−1^, we used a parallel-disk geometry with a gap width of 20 μm. It is a home-made narrow-gap device based on an MCR 501 rotational rheometer (Anton Paar, GmbH, Graz, Austria).^23–28^ To arrive at these low gap widths, the disks were replaced by glass plates (Edmund Optics) with an evenness of λ/4 (lower plate, 75 mm diameter) and λ/10 (upper plate, 50 mm diameter), where λ is the testing wavelength (633 nm). The plates were aligned with the help of actuators (Physik Instrumente (PI) GmbH & Co. KG, Karlsruhe Germany) and a confocal interferometric sensor (STIL, Aix-en-Provence, France) that also serves to independently adjust the gap width. By these means, the uncertainty in the gap width can be reduced to less than ±1 μm.^24^ The sketch of the setup is shown in Figure 1. The experiments were carried out at variations in gap widths below ±0.5 μm. Temperature was maintained at 25 ± 0.5 °C with a hood equipped with a temperature control system (H-Tronic TSM 125). Wet tissue inside the hood was used to minimize evaporation and maintain humidity. For further details on the setup and the alignment procedure, we refer to Dakhil and Wierschem, 2014.^24^

**Figure 1:**
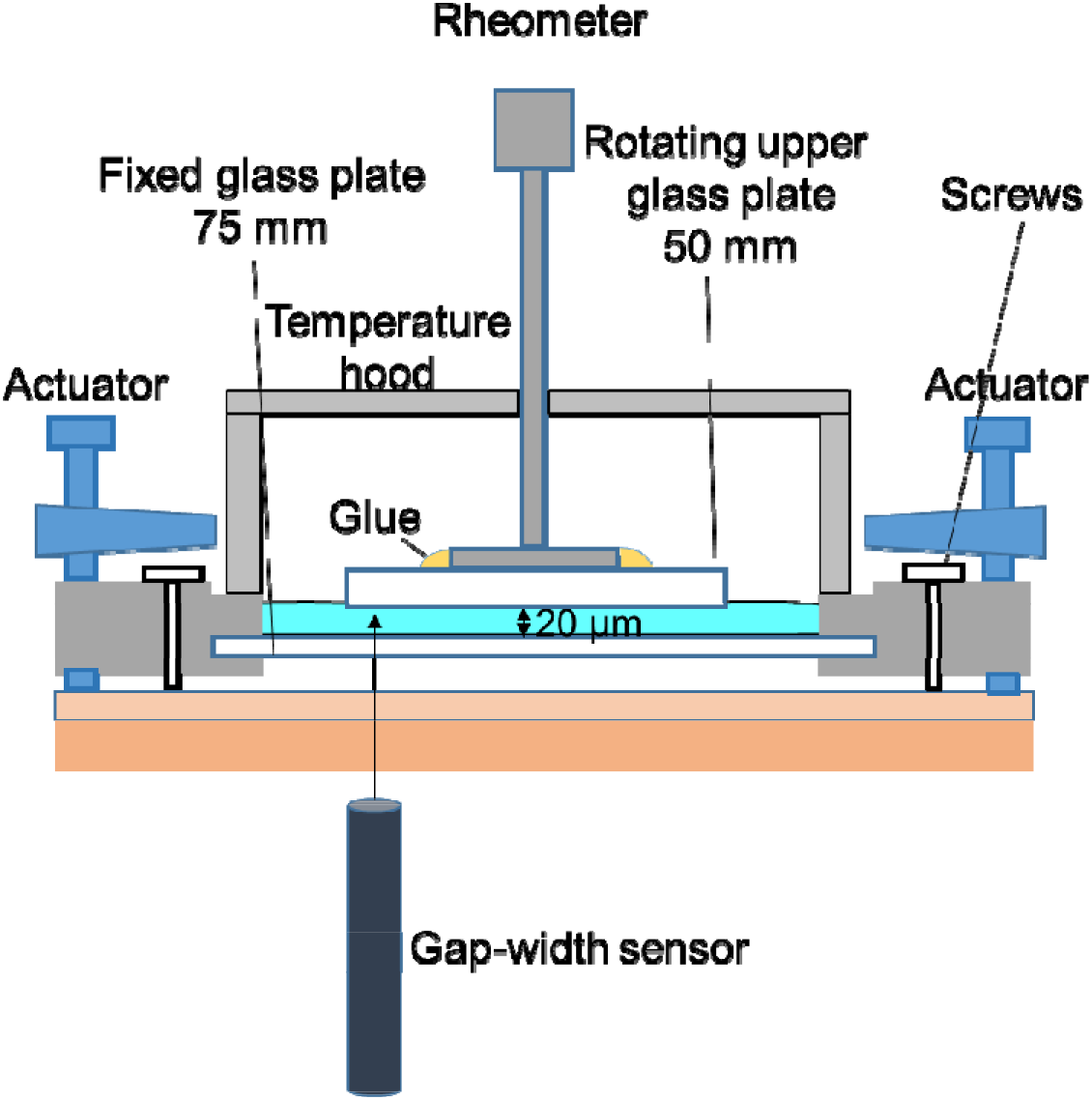
Illustration of the narrow-gap rheometer with homemade temperature control. The gap width between the rheometer plates was measured with a confocal interferometric sensor.

For shearing, we followed the established protocol from Dakhil et al., 2021.^23^ In the narrow-gap device, we pre-sheared the samples initially at a shear rate of 1,000 s^−1^ for 30 s and then again at 100 s^−1^ for 30 s followed by 5 min of rest. For each sample, measurements were carried out at 20 μm gap width along the shear rates from 1 s^−1^ – 150,000 s^−1^ with a time per measurement point of 10 s.

## 3 Results and discussion

### 3.1 Shear rheology of methyl cellulose solutions

Figure 2 shows the results from different devices for the viscosity and normal stress differences of the three solutions. The CC device yielded good results for the viscosity at low shear rates from 0.1 s^−1^ up to 20 s^−1^, 30 s^−1^ and 100 s^−1^ for 0.49% MC-PBS, 0.59% MC-PBS and 0.83% MC-PBS, respectively, but it cannot be used to measure normal stress differences. The oscillatory measurements show the complex viscosity results for angular frequencies between 0.1 s^−1^ and 80 s^−1^. The CP results cover a range of 100 – 25,000 s^−1^ and the PP results from 10 – 150,000 s^−1^. These devices were also used to determine the normal stress differences *N*_1_ and *N*_1_ – *N*_2_.

**Figure 2:**
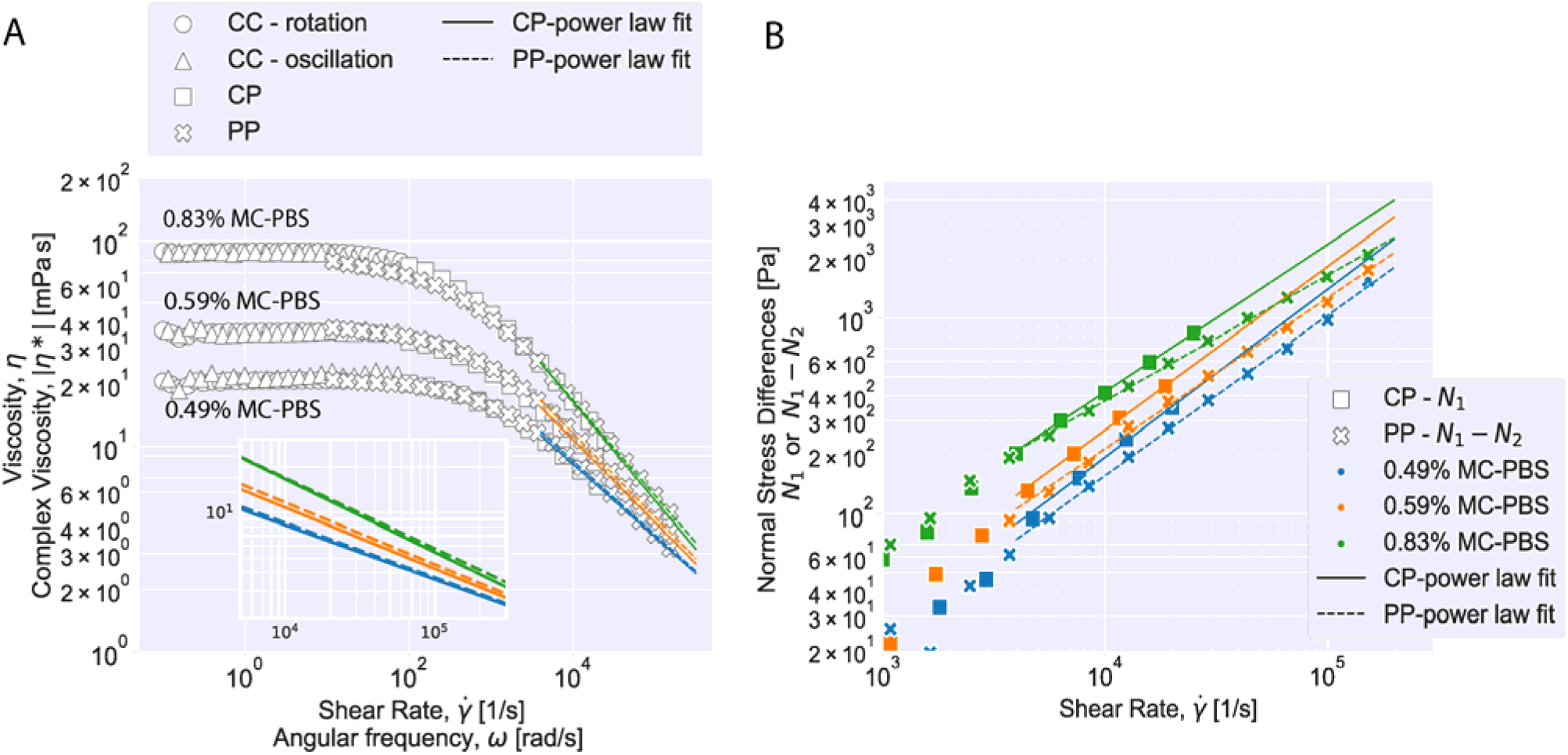
Shear and oscillation measurements of methylcellulose solutions with different measuring systems at 25 °C. Viscosity and normal stress differences fitted to a power law above 5,000 s^−1^. (A) Viscosity of methylcellulose solutions. The inset plot shows the fitted power law curves for the region, where the power law is valid. (B) Normal stress differences of methylcellulose solutions. 1^st^ normal stress difference with power law fit. (*N*_1_) is given for cone-plate (CP) and *N*_1_ - *N*_2_ is given for parallel disks (PP).

Figure 2A shows that the viscosity curves for all three solutions coincide well for the three different measurement systems. The CC results showed that the solutions reach a viscosity plateau at shear rates below 10 s^−1^. The CC results cover different shear rate regimes for each MC-PBS solution to avoid unreliable data points caused when reaching the shear rate limit of the CC device. Here, the measurements are affected by secondary flow instabilities, where eddies form at the bottom resulting in higher flow resistance.^29^ Thus, the viscosity curves from the CP and PP measurements are considered to be more accurate at shear rates above 100 s^−1^.

Measurements with CP and PP devices usually tend to show apparent yield stress at low shear rates (<100 s^−1^ for CP and <10 s^−1^ for PP). This is a known phenomenon for narrow gap rheometers and is likely caused by microscopic contaminations in the system or imperfect trimming.^24,29,30^ Thus, these shear rate regimes for CP and PP devices were not used for the data evaluation and the CC data is considered to deliver the most accurate results in this regime. The zero viscosity of the solutions was computed from the averaged CC data in the viscosity plateau up to 10 s^−1^. The results are given in Table 1.

**Table 1:**
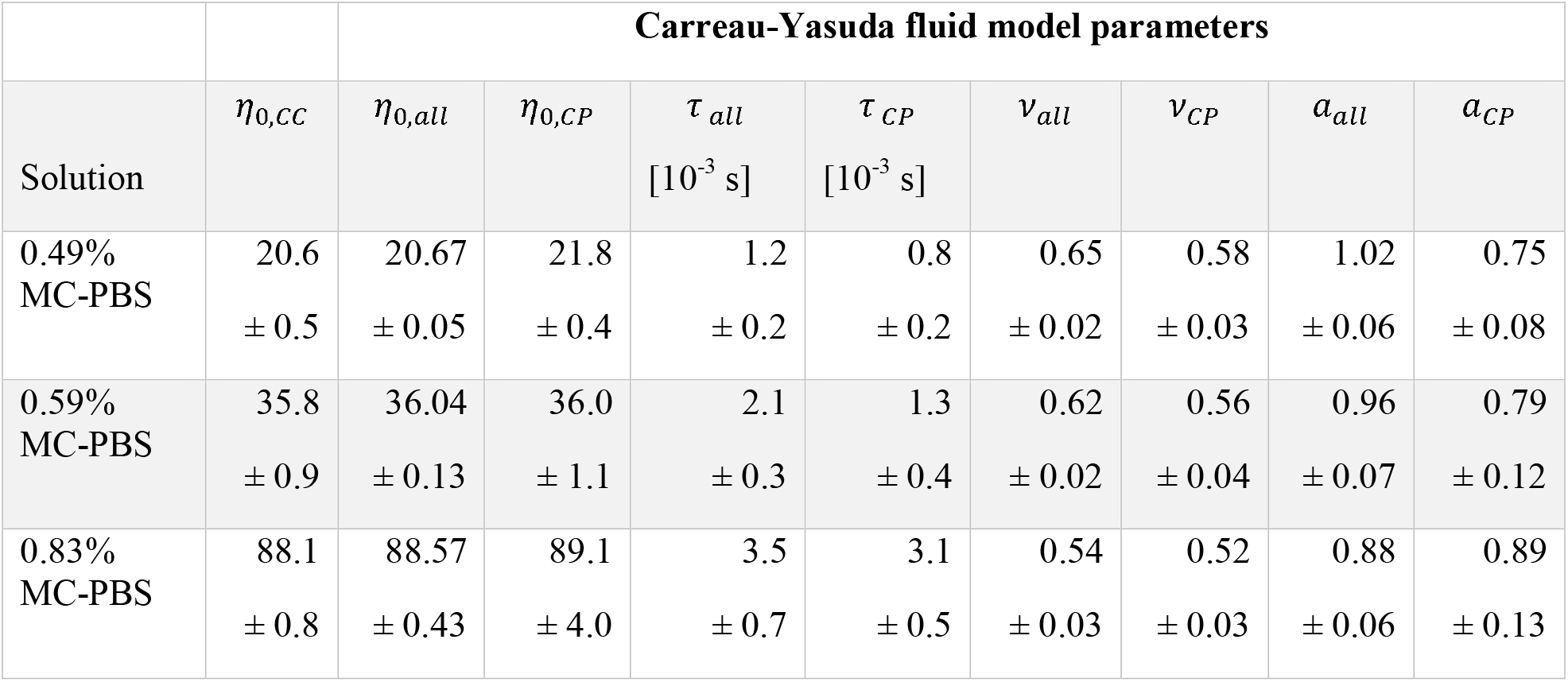
Representation of the experimental zero viscosities (*η*_0,*CC*_) from CC measuring system and Carreau-Yasuda fluid model parameters (*η*_∞_ = 0 mPa s) (value ± SD).

**Table 2:** Constitutive viscosity equations of MC-PBS solutions.

| Solution | Viscosity equation |
| --- | --- |
| 0.49% MC-PBS | $\eta(T, \dot{\gamma}) = 8.2 \cdot 10^{-7} \cdot e^{3691.8 \frac{[K]}{T}} \cdot \left(\frac{\dot{\gamma}}{\dot{\gamma}_0}\right)^{\left(0.0022 \frac{T}{[K]} - 0.99\right)}$ [Pa s] **(8)** |
| 0.59% MC-PBS | $\eta(T, \dot{\gamma}) = 1.5 \cdot 10^{-5} \cdot e^{3095.6 \frac{[K]}{T}} \cdot \left(\frac{\dot{\gamma}}{\dot{\gamma}_0}\right)^{\left(0.0024 \frac{T}{[K]} - 1.139\right)}$ [Pa s] **(9)** |
| 0.83% MC-PBS | $\eta(T, \dot{\gamma}) = 1.8 \cdot 10^{-5} \cdot e^{3351.6 \frac{[K]}{T}} \cdot \left(\frac{\dot{\gamma}}{\dot{\gamma}_0}\right)^{\left(0.0021 \frac{T}{[K]} - 1.12\right)}$ [Pa s] **(10)** |

For the oscillatory measurements with the CC device, the data points in the shear rate lower than 0.1 s^−1^and higher than 80 s^−1^ are not shown in Figure 2A due to exceeding the device torque limits and instrument inertia. In the reliable shear rate range, the measured complex viscosity and shear viscosity values match very well, fulfilling the Cox-Merz rule (Eq. (3)):

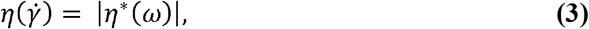

where *η*\* represents the absolute complex viscosity and *ω* the angular frequency.

For shear rates over 1,000 s^−1^, CP and PP measurements gave very similar viscosity curves. The solutions change from the viscosity plateau at lower shear rates into a power law behavior at higher shear rates, which can be considered fully developed above 5,000 s^−1^ for all solutions. The shear thinning will start at lower shear rates for higher MC concentrations, which is typical for polymer solutions. The CP and PP viscosity curve were analyzed for shear rates > 5,000 s^−1^ with a power law given by the following equation:

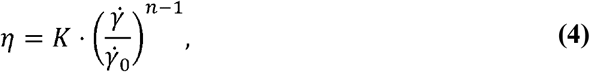

where 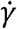 is the shear rate, 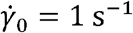, *K* is the *flow consistency index* and *n* the *flow behavior index*. The resulting fit values for *K* and *n* are reported in Table S1. It can be seen that values for *n* from PP and CP are similar for two digits within the error margin, which tells that both devices predict a similar shear thinning behavior for increasing shear rates. Values for *K* are close as well but show larger deviation, which can be attributed to a slight offset of the absolute viscosity between the two devices. The nice agreement between CP and PP measurements together with the fact that the samples obey the power law shows that measurements with the CP up to 25,000 s^−1^can be used to extrapolate the viscosities to the highest shear rates studied with the PP i.e. 150,000 s^−1^. The results from both devices show that *K* increases with the MC concentration while *n* decreases, which means that the viscosity increases and the solutions show stronger shear thinning.

Combining all reliable data points measured with the three devices in rotation yields an accurate viscosity curve that covers the full range of shear rates investigated here. The viscosities were modelled with the Carreau-Yasuda model given in equation (5):

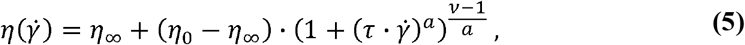

with the infinite-shear viscosity *η*_∞_, the zero viscosity *η*_0_, the relaxation time *τ*, the Yasuda model parameter *a* and the power law index *ν*. We performed model fits on the combined curves from all three devices and on the curves from CP results only, to check if the range covered by the CP device is enough to extract an accurate model curve. Since the shear rates used for this study were not high enough for deviations from the power law into the infinite-shear viscosity plateau, we fixed *η*_∞_ = 0, which is a common convention for viscosity curves. The results are plotted in Figure 3 and the fit parameters are given in Table 1.

**Figure 3:**
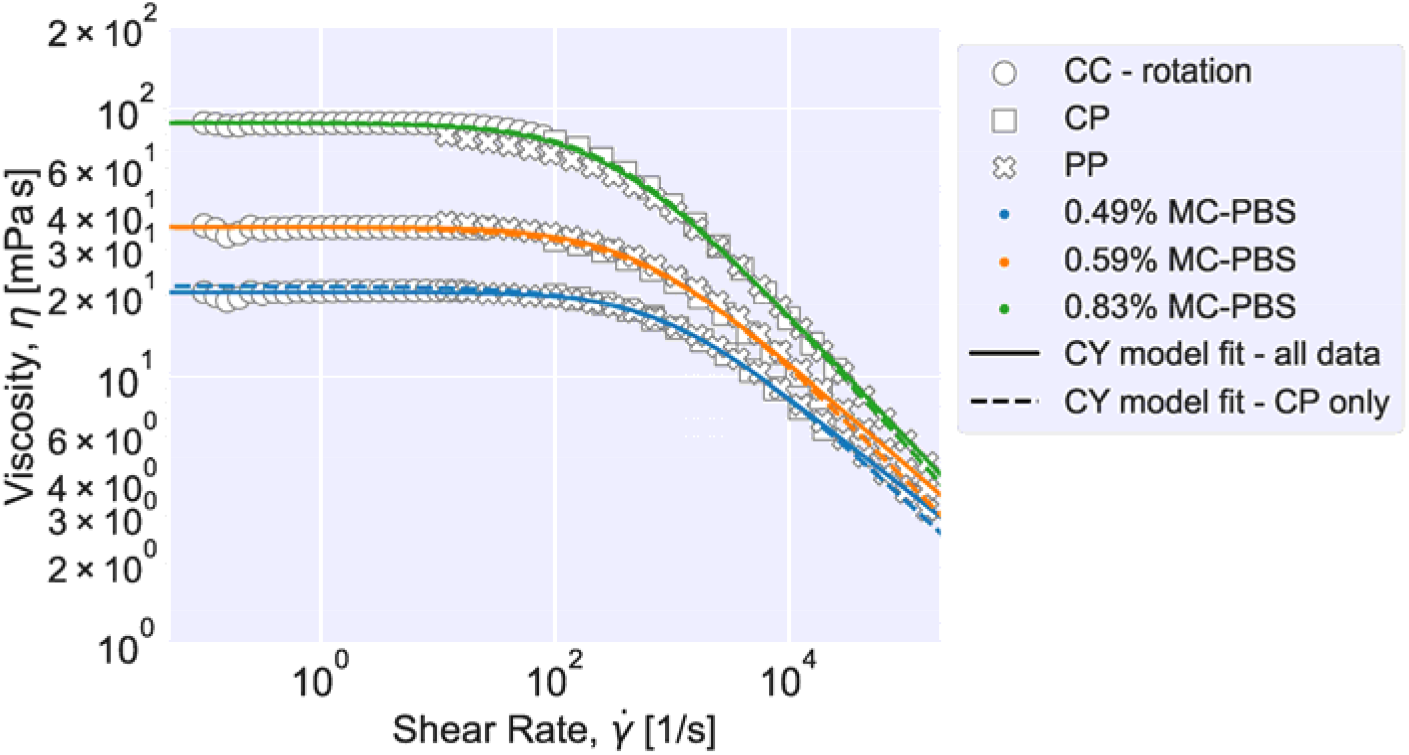
Carreau-Yasuda (CY) model fit to data of rotational shear viscosity. Results from rotational CC, CP and PP measurements for all three solutions and the corresponding fitted functions using the CY model on all data points (solid line) and when only using the CP data points (dashed line).

It can be seen that the Carreau-Yasuda model is well suited to describe the full viscosity curve of all three solutions used for this study. The values for the zero viscosity from fitting all data points coincide well with the fit for the CP-only data. Both values are close to the zero viscosity values extracted from the mean viscosity computed from the CC results below 10 s^−1^ (*η*_0,*cc*_, see Table 1). The relaxation time increases with increasing MC concentration and is higher for all fits for the complete curve, compared to the CP-only data. The relaxation time is connected to the shear rate, at which the polymer solution changes from Newtonian to power law behavior: 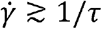. Also, the power law index is greater for the fit on the full curve, which means that including all data points leads to a prediction of stronger shear thinning at high shear rates. The predicted power law indices are close to the values computed from the pure power law fitted on the data at high shear rates only (see equation (4) and Table S1). The Yasuda model parameter *a* correlates with the length of the intermediate region from Newtonian to power law behavior. Within fitting uncertainty, it is about constant.

Normal stress differences (*N*_1_ and *N*_1_ – *N*_2_) were derived from CP and PP measurements. While data below 1,000 s^−1^ is dominated by noise, the data for larger shear rates shows a power law relation between *N*_1_ and the shear rate. Like for the viscosity data, we fitted a power law to the *N*_1_ data for shear rates above 5,000 s^−1^ with the following equations^31^:

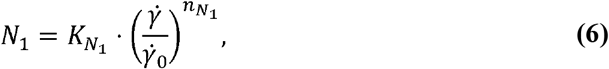

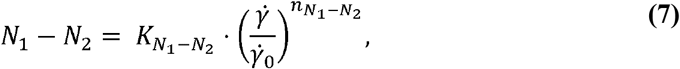

where 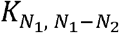 is the *stress consistency index* and 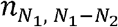 the *stress behavior index*. The fit parameters are listed in Table S1. The results show that the stress differences increased with buffer concentration and so did 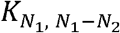 while 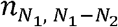 decreased.

### 3.2 Temperature dependence

The viscosity functions at different temperatures can be superposed to a master curve by time-temperature superposition.^32,33^ To determine the shift factor, *a_r_*, with respect to the viscosity function at the reference temperature, the viscosity functions were first fitted to a representative function. The shift factor is given by 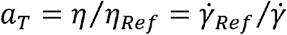, where the index *Ref* refers to the reference temperature, which was chosen to be 25 °C. To superpose the master curves, we define the reduced shear rate 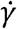 · *a_T_* and the reduced viscosity *η*/*a_T_*. The master curves for the different concentrations are shown in Figure 4A and the corresponding shift factors in Figure 4B.

**Figure 4:**
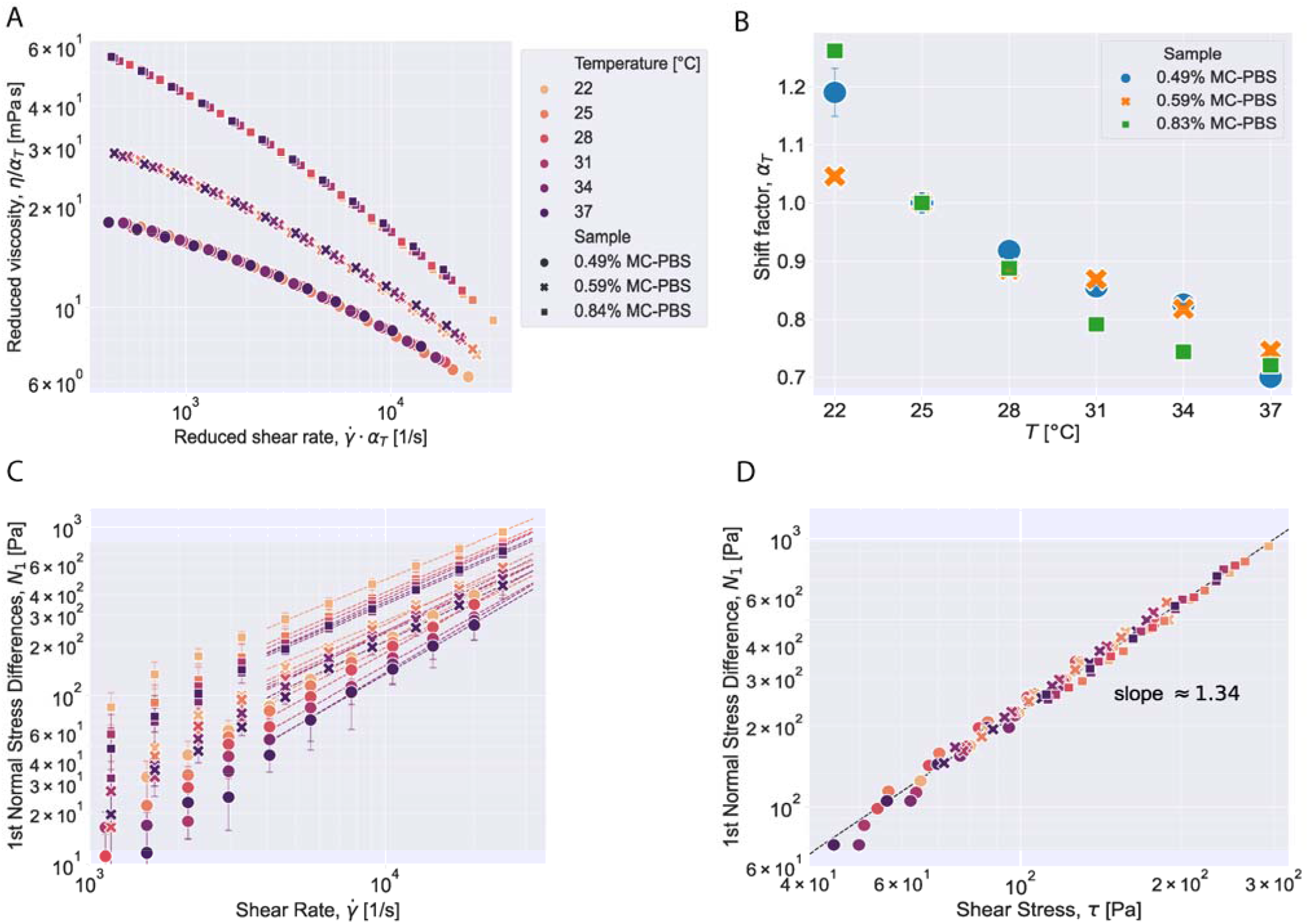
Temperature dependent shear rheology of methyl cellulose. (A) Master curves of MC-PBS solutions by time-temperature superposition and (B) corresponding shift factors for master curves. Error bars show the fitting uncertainty. (C) Temperature dependence of 1^st^ normal stress differences for all MC-PBS solutions. Error bars represent the standard deviation of triple measurements and dashed lines show the power law fit. (D) 1^st^ normal stress difference over shear stress for different solutions and temperatures fitted to a linear line.

As seen in Figure 4A, the viscosity functions at different temperatures superpose nicely to a master curve. The fitting uncertainty for the shift factors is shown as error bars in Figure 4B. The temperature dependence of the shift factor for the medium concentration follows a linear trend. The shift factors for the lowest concentration coincide with those at medium concentration except for the lowest temperature studied. The shift factors for the highest concentration deviate from this trend. This might be due to proximity to thermo-rheologically complex regimes such as gelation.^32^ Indeed, MC solutions may gel at higher concentrations when heated. Added salt lowers the gelation temperature.^34^ Xu *et al*. observed, for instance, that 0.8 M sodium chloride can shift in the gelation temperature of an aqueous 0.93 wt.% MC solution by about 20 °C to 40 °C. Although the PBS-solution used is about 0.15 M sodium chloride solution, it cannot be ruled out that the proximity of the gel point may yield deviations from the linear trend.

Shear viscosity and *N*_1_ decrease with increasing temperature at all shear rates for all three solutions. Power law behavior was observed for the first normal stress difference for shear rates over 5,000 s^−1^ at all temperatures and the power law parameters 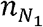 and 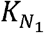 were determined by fitting equation (4) and (6) and are reported in Table S2. Plotting the data for *N*_1_ over the shear stress for all temperatures and solutions (Figure 4D) collapses all data points on one line with slope 1.34 in the double-logarithmic plot, independent of temperature or MC concentration. This behavior was described for other polymer solutions before.^31,35,36^

At all temperatures studied, the viscosity curves followed a power law according to equation (4) at shear rates greater 5,000 s^−1^ (see fig. S1). For this regime, we formulated temperature dependencies of the power law parameters *K* and *n* (see SI text and fig. S1). This yielded constitutive viscosity equations for each solution dependent on shear rate and temperature. These are given in **Error! Reference source not found**.:

### 3.3 Concentration dependence

To get a better picture how the MC concentration influences the overall shear rheology of the MC-PBS solutions, we measured solutions with different MC concentrations in the CP device. Figure 5A shows the viscosity curves of eight different MC solutions with MC concentrations between 0.3 – 1.0 w/w% at 25 °C. Lower concentrations were below the low torque limit of the CP device. The Carreau-Yasuda model (Eq. (5)) was used to represent the viscosity behavior of the solutions in the shear rate range between 100 s^−1^ – 25,000 s^−1^ for solutions < 0.8% MC and between 10 s^−1^ – 25,000 s^−1^ for solutions ≥ 0.8% MC. The model parameters estimated from the Carreau-Yasuda model are given in Table S4.

**Figure 5:**
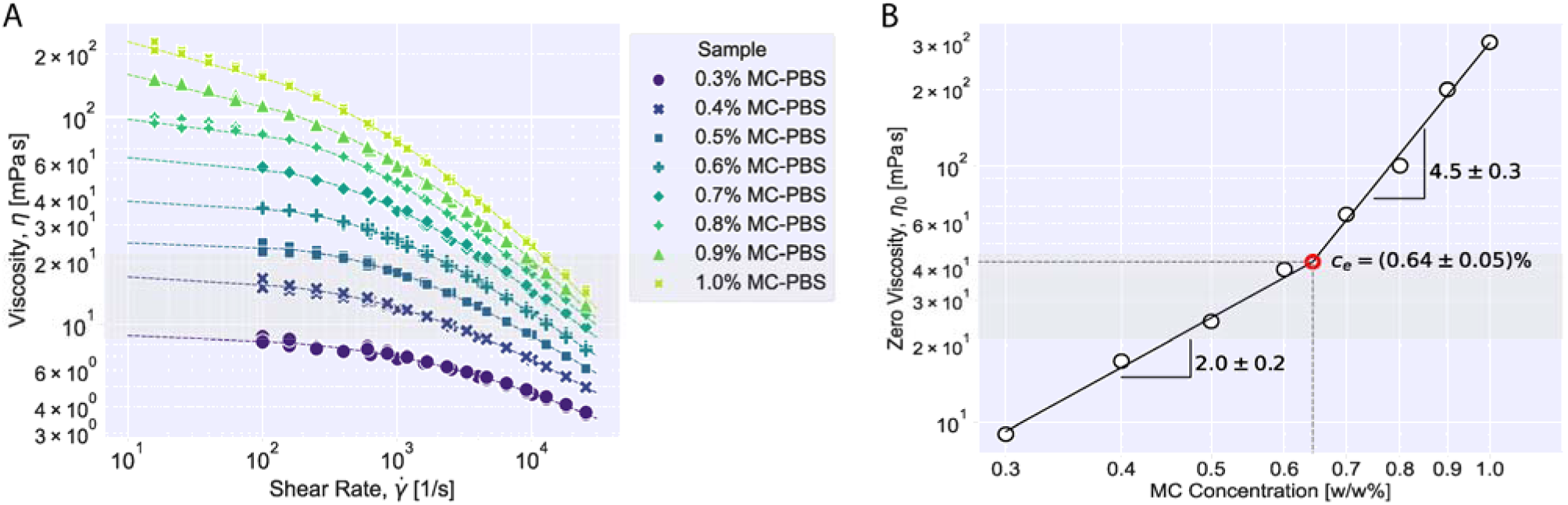
Concentration dependent shear rheology at 25 °C. (A) Viscosity of different concentrated MC-PBS solutions in the shear rate range between 10 s^−1^ and 25,000 s^−1^. The dashed lines represent the fitting to CY model. (B) Representation of zero viscosity depending on MC-PBS concentration. *c_e_* is given for the entanglement concentration of methylcellulose at 25 °C.

Figure 5B shows the zero-viscosity change depending on MC concentration. Power law dependency of zero viscosity on concentration is shown by the dashed lines based on the relation:

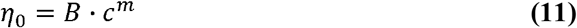

where *η*_0_ is zero viscosity, *B* represents a pre-factor, *c* is concentration in (w/w%) and is m the slope in the double-logarithmic plot. The results for the MC solutions have shown that there is an apparent slope change from approximately 2.0 to 4.5 at a concentration of 0.64 ± 0.05 (w/w%). This transition point is defined as *entanglement concentration* (*c_e_*) and can be explained as the transition of MC-PBS polymer solution from the semidilute unentangled to the semidilute entangled regime. A slope of 2±0.2 for the semi-dilute unentangled regime and a value of 4.5±0.3 for the semidilute entangled regime are within error margins in agreement with scaling predictions for neutral polymer solutions in a theta solvent, which is a valid approximation for the solutions used here.^37,38^

The 1^st^ normal stress differences were also determined for the concentration study and the results can be found in Figure S2. The figure shows the power law behavior of *N*_1_ (Eq. (6)) at all concentrations at shear rates higher than 5,000 s^−1^. *N*_1_ values increased with the increasing concentration.

### 3.4 Discussion

#### 3.4.1 Rheology of the solutions

We characterized the shear rheology of three MC-PBS solutions used for cell mechanical measurements in real-time deformability cytometry (RT-DC) by combining the results from three different rheometer setups: concentric-cylinders (CC), cone-plate (CP) in counter-rotation and narrow-gap parallel-disks (PP) to cover a wide range of shear rates from 0.1 – 150,000 s^−1^. The results from the three techniques were in good agreement with each other and showed that the viscosity of the solutions followed a Carreau-Yasuda model over the full range of measured shear rates. For shear rates below 100 s^−1^, the solutions were measured in rotation and oscillation and it was observed that the solutions follow the Cox-Merz-relation.

Furthermore, we showed that it was sufficient to measure the rheology of the buffers in the intermediate shear rate regime of 100 – 25,000 s^−1^ in the commercially available CP configuration and then extrapolate the viscosity curve using the Carreau-Yasuda model or a power law model for higher shear rates.

At shear rates over 1,000 s^−1^, all solutions exhibited normal stress differences, which dominated the shear stresses at high shear rates. This is an expected behavior for polymer solutions and indicates the viscoelastic nature of the MC-PBS solutions.^31^ In neutral polymer solutions like in the studied system, normal-stress differences occur mainly due to the deformation and orientation of the polymers. This influence is usually disregarded when estimating the stresses in microfluidic configurations when using MC.^9,11,12,17,18^

Probing the solutions at temperatures between 22 – 37 °C, which is a typical temperature range for biological samples, showed that the viscosity functions at different temperatures could be superposed to a master curve by time-temperature superposition.^32,33^ The knowledge about the temperature behavior of the shift factors allows to extrapolate the viscosity range beyond the shear rate range accessible by the CP device. For the power-law behavior that the solutions exhibit for viscosity and normal stress differences at shear rates beyond 5,000 s^−1^, we used the results to derive an empirical viscosity equation for the three solutions depending on shear rate and temperature. These equations can be used to compute the viscosities in the entire shear-rate and temperature range studied. The parameters that describe the influence of the temperature on the viscosity showed no trend for different MC concentrations. This result can be used to derive the temperature dependence for solutions with any MC concentration.

To determine the entanglement concentration of the MC-PBS solutions, we measured solutions with MC concentration from 0.3 – 1.0 w/w%. The Carreau-Yasuda model fits the viscosity data well in this range and could be used to estimate the zero-viscosities *η*_0_. The first normal stress difference followed a power law for all concentrations. From the relation of *η*_0_ to the MC concentration, we found that the entanglement concentration is at about 0.6 w/w%.

#### 3.4.2 Implications for microfluidic experiments

Microfluidic configurations often use channel geometries with a rectangular cross section because such channels can be easily produced by using standard soft lithography methods. In contrast to the rheometers used for this study, in pressure-driven flows such as in microfluidic channels, liquids do not experience a uniform shear rate. Shear stresses are highest at the channel walls and zero at the channel centerline. This results in viscosity and accordingly shear-rate distributions and normal-stress differences within the channel. For Newtonian liquids, the velocity profile, and with that the distribution of shear rates, is independent of the material properties and can be determined by knowing only the flow rate inside the channel. For a shear thinning liquid, the velocity profile is flatter in the central part and steeper at the walls. For a power-law liquid, it can be expressed by the flow behavior index *n*. Since there is a continuous shear-rate distribution in the microfluidic channel for a given flow condition, the MC solutions will have an according viscosity distribution within the channel cross section. To compute the stiffness of cells from RT-DC experiments, the cell deformation is mapped to a Young’s modulus.^17^ The resulting Young’s modulus scales linearly with the liquid’s viscosity. To define this viscosity value, Herold^15^ proposed to use the viscosity at the *wall shear rate 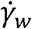*. That this is a valid approach was recently confirmed by Wittwer et al. by employing finite element simulations to mimic the conditions in RT-DC experiments.^39^ Son derived a formula to determine the wall shear rate in a rectangular channel for a power law liquid at a given flow rate Q:^40^

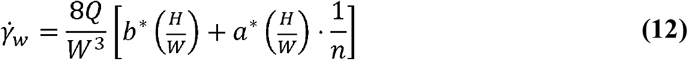

where *W* is the channel width, *H* is the channel height and *H* < *W*, *a*\* and *b*\* are geometrical factors dependent on the aspect ratio of the channel *H*/*W* and were determined numerically (for a list of values, see Son^40^) and *n* is the flow behavior index. By inserting equation (12) into equation (4), one can calculate the *effective viscosity* inside the channel. In the case of RT-DC channels, *H* = *W* and *a*\* = 0.2121 and *b*\* = 0.6771. This leads to the following equation to calculate the effective viscosity in RT-DC:

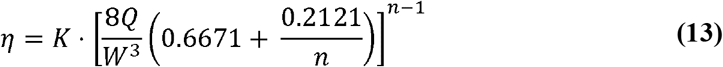

with the flow consistency index *K* and the flow behavior index *n*.

It is known that the viscoelasticity of fluids influences the particle migration in channel flow.^41^ In shear thinning liquids, particle migration towards the channel wall was observed^42^ while in the elasticity-dominated regime, particles tend to focus along the centerline.^41^ This indicates that forces act transversely to the flow direction in the channel flow of viscoelastic liquids. Particle tracing experiments could reveal how these normal stresses act in the channel flow of the studied MC solutions.

In the case of cell deformability experiments this would lead to additional normal stresses on the cell surface. In solutions with larger relaxation times, the slower dynamics may affect cell measurements as, for instance, a subsequent cell may be affected by the incomplete relaxation caused by its forerunner. Furthermore, if the measurement happens at a timescale in the range of the relaxation times of the MC solutions, it will influence the effective viscosity inside the channel and subsequently the stresses on the cell. Future investigations in this direction are needed to quantify the magnitude of these stresses and how they influence the cell shape.

## 4 Conclusions

With this study, we aimed to provide a full shear rheological description of MC-PBS solutions used in RT-DC and to better understand the behavior of MC-PBS solutions dependent on temperature and MC concentration. These results will be useful to calculate stresses exerted by the solutions during flow in microfluidic systems.^8^ To our knowledge, this is the first study to describe normal stress differences exerted by MC solutions at high shear rates, which highlight the viscoelastic nature of the solutions. These effects should be accounted for when estimating stresses caused by the solutions in microfluidic experiments, also in extensional flows.

## 5 Author Contributions

Conceptualization: A.W., J.G., and F.R.; Experiments CC-device: M.N.; Experiments CP-device: B.B. and F.R.; Experiments PP-device: S.K.B.; Writing – original draft: B.B. and F.R., Writing – editing and review: all authors.

## 7 Acknowledgments

We would like to thank Cornelia Liebers for help with the production of MC-PBS solutions. We would also thank Dr. José Alberto Rodríguez Agudo from Anton Paar GmbH for advice on operation of the CP device. The authors acknowledge the financial support through the base funding of the Max Planck Society to JG.

## 8 Data Availability Statement

The datasets and analysis scripts for this study are available on gitlab: https://gitlab.gwdg.de/shear-rheology-of-methyl-cellulose-solutions (persistent identifier (PID): 21.11101/0000-0007-F9A5-6; see https://www.pidconsortium.net)

## 9 Supplementary Material

Supplementary material is provided in an additional file.

