## Supplementary Information for "Shear rheology of methyl cellulose based solutions for cell mechanical measurements at high shear rates"

### 1 Temperature dependence of power law parameters

To characterize the effect of temperature on the solutions, we investigated the temperature dependency of the fitting parameters $n$ and $K$. It is shown in Figure S1. Here, we assumed an empirically derived linear relationship between $n$ and $T$ and an exponential relation between $K$ and $1/T$ motivated by the Arrhenius law:

$$n=\alpha\cdot T+\beta$$

(1)

$$K=A\cdot e^{\frac{\lambda}{T}}$$

(2)

The corresponding fitting parameters were also determined and noted as $\alpha$ and $\beta$ for linear dependency of $n$, $A$ and $\lambda$ for the exponential dependency of $K$. These parameters are listed in Table S2, depending on the MC concentration. It can be seen that the flow behavior index $n$ increases with temperature, which means that shear thinning is less pronounced at higher temperatures. The flow consistency index $K$ shows an exponential increase with $1/T$, which corresponds to the Arrhenius law for the viscosity of liquids.

For the viscosity, it can be seen that the parameter $\alpha$ did not strongly depend on the MC concentration. $\alpha$ describes how strongly the flow behavior index $n$ correlates with the temperature and, hence, how shear thinning is affected by temperature. Since $\alpha\cdot T$ is small compared to $\beta$ for our temperature range, the shear thinning exponent depended only weakly on temperature and $\alpha$ was independent of the MC concentration for the solutions investigated here. The parameter $\lambda$ also stayed constant, within error margins, for all three MC concentrations and the temperature behavior was mainly determined by the pre-factor $A$. $\lambda$ is related to the activation energy of MC. These results show that $\alpha$ and $\lambda$ are material constants for the MC-PBS solutions, which can be used to describe the temperature dependence at any MC concentration.

The temperature dependence of the power law fit parameters for the first normal stress difference, $K_{N_{1}}$ and $n_{N_{1}}$, are depicted in Figure S1B and it can be seen that there is no strong correlation with the temperature, here.

Knowing all of the parameters affecting the rheological behavior of methyl cellulose solutions helps us to form a constitutive equation for each solution, which are valid for shear rates beyond 5,000 s^-1^. These equations are listed in main text Table 2 and allow obtaining the viscosity of these solutions at a certain temperature and shear rate that are relevant for measuring biological cells in microfluidic applications.

### 2 Supplementary figures

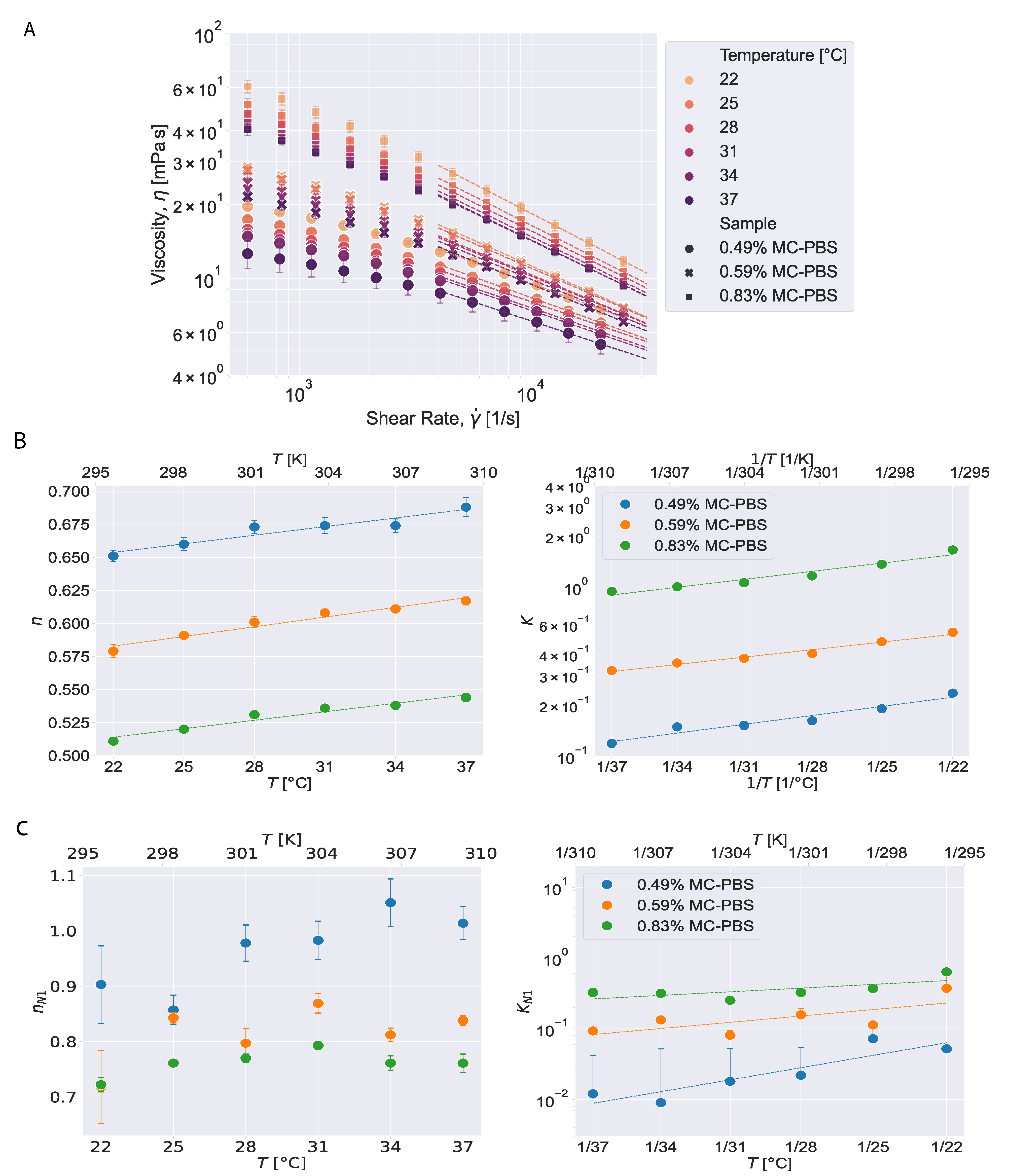

Figure S1: Temperature dependence of viscosity and 1^st^ normal stress differences. (A) Viscosity curves of all MC-PBS solutions with power law fit at shear rates higher than 5,000 s^-1^. (B) Temperature dependency of $\boldsymbol{n}$ and $\boldsymbol{K}$. (C) Temperature dependency of $\boldsymbol{n}_{\boldsymbol{N}\boldsymbol{1}}$ and $\boldsymbol{K}_{\boldsymbol{N}\boldsymbol{1}}$ (mean ± SD).

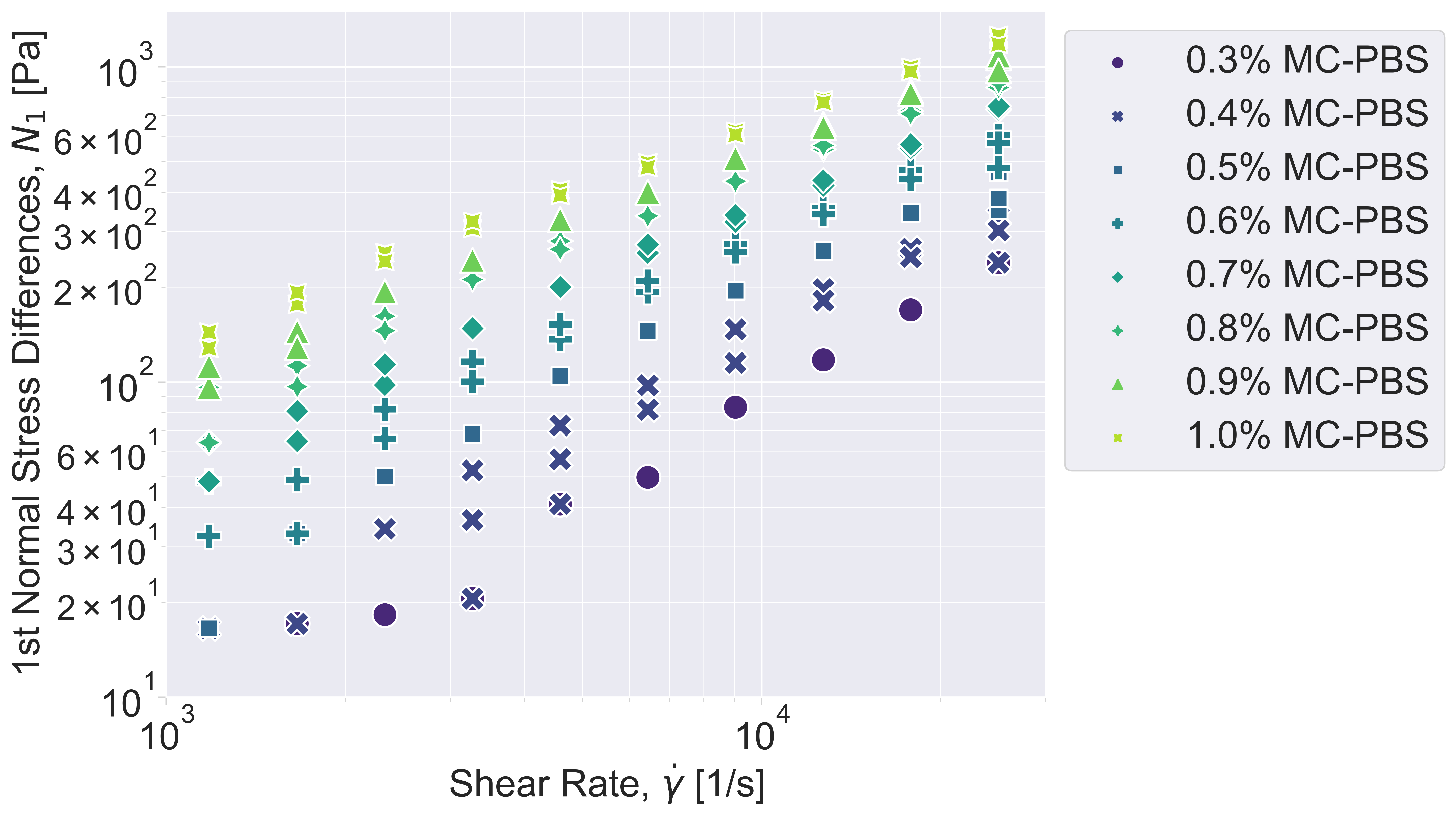

Figure S2: 1^st^ normal stress differences for solutions with varying MC concentration.

### 3 Supplementary tables

Table S1: Comparison of power law fit parameters for different devices (value ± SD)

| **Viscosity – shear rate** | | | |
| --- | --- | --- | --- |
|  | Solution | Device Type | |
|  |  | cone-plate | plate-plate |
| $n$ | 0.49% MC-PBS | 0.645 ± 0.007 | 0.641 ± 0.002 |
|  | 0.59% MC-PBS | 0.598 ± 0.004 | 0.593 ± 0.003 |
|  | 0.83% MC-PBS | 0.520 ± 0.002 | 0.536 ± 0.001 |
| $K$ [Pa s] | 0.49% MC-PBS | 0.21 ± 0.06 | 0.23 ± 0.03 |
|  | 0.59% MC-PBS | 0.43 ± 0.04 | 0.49 ± 0.03 |
|  | 0.83% MC-PBS | 1.35 ± 0.02 | 1.19 ± 0.01 |
| **Normal stress differences (**${\boldsymbol{N}_{\boldsymbol{1}}\boldsymbol{or N}}_{\boldsymbol{1}}\boldsymbol{-}\boldsymbol{N}_{\boldsymbol{2}}$**) – shear rate** | | | |
| $n_{N_{1}, N_{1}-N_{2}}$ | 0.49% MC-PBS | 0.85 ± 0.06 | 0.818 ± 0.014 |
|  | 0.59% MC-PBS | 0.83 ± 0.03 | 0.770 ± 0.016 |
|  | 0.83% MC-PBS | 0.75 ± 0.02 | 0.641 ± 0.008 |
| $K_{N_{1}, N_{1}-N_{2}}$ [Pa s] | 0.49% MC-PBS | 0.07 ± 0.03 | 0.08 ± 0.01 |
|  | 0.59% MC-PBS | 0.12 ± 0.03 | 0.18 ± 0.03 |
|  | 0.83% MC-PBS | 0.40 ± 0.04 | 1.02 ± 0.08 |

Table S2: Fit parameters of temperature dependent study (value ± SD)

|  | **Viscosity – shear rate** | | | | | | |
| --- | --- | --- | --- | --- | --- | --- | --- |
|  | **Solution** | **Temperature [°C]** | | | | | |
|  |  | **22** | **25** | **28** | **31** | **34** | **37** |
| $\boldsymbol{n}$ | 0.49% MC-PBS | 0.651  $\pm$ 0.004 | 0.660  $\pm$ 0.005 | 0.673  $\pm$ 0.005 | 0.674  $\pm$ 0.006 | 0.674  $\pm$ 0.005 | 0.688  $\pm$ 0.007 |
|  | 0.59% MC-PBS | 0.579  $\pm$0.005 | 0.591  $\pm$ 0.002 | 0.601  $\pm$ 0.004 | 0.608  $\pm$ 0.003 | 0.611  $\pm$ 0.003 | 0.617  $\pm$ 0.003 |
|  | 0.83% MC-PBS | 0.511  $\pm$ 0.002 | 0.520  $\pm$ 0.001 | 0.531  $\pm$ 0.002 | 0.536  $\pm$ 0.002 | 0.538  $\pm$ 0.003 | 0.544  $\pm$ 0.003 |
| $\boldsymbol{K}$ **[Pa s]** | 0.49% MC-PBS | 0.236  $\pm$ 0.010 | 0.191  $\pm$ 0.009 | 0.162  $\pm$0.008 | 0.152  $\pm$ 0.009 | 0.149  $\pm$0.007 | 0.119  $\pm$ 0.007 |
|  | 0.59% MC-PBS | 0.541  $\pm$ 0.027 | 0.476  $\pm$ 0.010 | 0.405  $\pm$ 0.016 | 0.379  $\pm$ 0.009 | 0.356  $\pm$ 0.011 | 0.321  $\pm$ 0.008 |
|  | 0.83% MC-PBS | 1.662  $\pm$ 0.028 | 1.367  $\pm$ 0.019 | 1.167  $\pm$ 0.021 | 1.065  $\pm$ 0.024 | 1.006  $\pm$ 0.032 | 0.945  $\pm$ 0.029 |
|  | **1^st^ normal stress difference (**$N_{1}$**) – shear rate** | | | | | | |
| $\boldsymbol{n}_{N_{1}}$ | 0.49% MC-PBS | 0.903  $\pm$ 0.007 | 0.857  $\pm$ 0.027 | 0.978  $\pm$ 0.033 | 0.983  $\pm$0.034 | 1.051  $\pm$ 0.043 | 1.014  $\pm$ 0.030 |
|  | 0.59% MC-PBS | 0.718  $\pm$ 0.066 | 0.843  $\pm$0.009 | 0.797  $\pm$ 0.026 | 0.869  $\pm$ 0.017 | 0.812  $\pm$ 0.012 | 0.838  $\pm$ 0.008 |
|  | 0.83% MC-PBS | 0.722  $\pm$ 0.013 | 0.761  $\pm$ 0.004 | 0.770  $\pm$ 0.007 | 0.793  $\pm$ 0.007 | 0.761  $\pm$ 0.013 | 0.761  $\pm$ 0.017 |
| $\boldsymbol{K}_{N_{1}}$ **[Pa s]** | 0.49% MC-PBS | 0.052  $\pm$ 0.003 | 0.072  $\pm$ 0.018 | 0.022  $\pm$ 0.007 | 0.018  $\pm$ 0.006 | 0.009  $\pm$ 0.003 | 0.012  $\pm$ 0.003 |
|  | 0.59% MC-PBS | 0.373  $\pm$0.232 | 0.113  $\pm$ 0.010 | 0.157  $\pm$0.039 | 0.081  $\pm$ 0.014 | 0.133  $\pm$ 0.016 | 0.093  $\pm$ 0.007 |
|  | 0.83% MC-PBS | 0.635  $\pm$0.076 | 0.370  $\pm$ 0.014 | 0.325  $\pm$ 0.022 | 0.252  $\pm$ 0.017 | 0.315  $\pm$ 0.039 | 0.323  $\pm$ 0.051 |

Table S3: Fitting parameters for $\boldsymbol{n}$ and $\boldsymbol{K}$ dependent on temperature (fit value ± SD)

| **Solution** | $\boldsymbol{\alpha}$ **[**$\boldsymbol{1}/\mathbf{K}$**]** | $\boldsymbol{\beta}$ | $\boldsymbol{A}$ **[Pa s]** | $\boldsymbol{\lambda}$ **[**$\mathbf{K}$**]** |
| --- | --- | --- | --- | --- |
| **0.49% MC-PBS** | 0.0022  $\pm$ 0.0004 | 0.01  $\pm$ 0.11 | $0.8\cdot{10}^{-6}$  $\pm1.0\cdot{10}^{-6}$ | 3691.8  $\pm$475.2 |
| **0.59% MC-PBS** | 0.0024  $\pm$ 0.0003 | -0.14  $\pm$ 0.08 | $1.5\cdot{10}^{-5}$  $\pm1.2\cdot{10}^{-5}$ | 3095.6  $\pm$ 242.5 |
| **0.83% MC-PBS** | 0.0021  $\pm$ 0.0003 | -0.12  $\pm$ 0.08 | $1.8\cdot{10}^{-5}$  $\pm2.5\cdot{10}^{-5}$ | 3351.6  $\pm$ 412.4 |

Table S4: List of zero viscosity and Carreau-Yasuda model parameters of different methylcellulose solutions (value ± SD)

| Solution | $\boldsymbol{\eta}_{\boldsymbol{0}}$ [mPa s] | $\boldsymbol{\tau}$  [10^-3^ s] | $\boldsymbol{\nu}$ | $\boldsymbol{a}$ |
| --- | --- | --- | --- | --- |
| 0.3% MC-PBS | 9.00 $\pm$ 0.48 | 0.35 $\pm$ 0.53 | $0.65\pm$0.14 | 0.58 $\pm$ 0.20 |
| 0.4% MC-PBS | 17.36 $\pm$0.68 | 0.80 $\pm$ 0.42 | 0.61 $\pm$ 0.06 | 0.66 $\pm$ 0.12 |
| 0.5% MC-PBS | 24.79 $\pm$ 0.52 | 1.34 $\pm$ 0.23 | 0.61 $\pm$0.02 | 0.99 $\pm$ 0.13 |
| 0.6% MC-PBS | 39.61 $\pm$ 0.83 | 1.54 $\pm$ 0.21 | 0.55 $\pm$ 0.02 | 0.91 $\pm$ 0.09 |
| 0.7% MC-PBS | 67.92 $\pm$2.46 | 1.90 $\pm$ 0.40 | 0.51 $\pm$ 0.04 | 0.82 $\pm$ 0.11 |
| 0.8% MC-PBS | 100.66$\pm$ 1.61 | 2.43 $\pm$ 0.45 | 0.46 $\pm$ 0.03 | 0.79 $\pm$ 0.06 |
| 0.9% MC-PBS | 199.77 $\pm$ 5.76 | 1.01$\pm$ 0.42 | 0.25$\pm$ 0.06 | 0.43 $\pm$ 0.03 |
| 1.0% MC-PBS | 305.36 $\pm$ 19.49 | 0.97 $\pm$0.85 | 0.17 $\pm$0.14 | 0.41 $\pm$0.05 |
